# *Cypripedium wardii* (Orchidaceae) employs pseudopollen with both reward and deception to attract both flis and bees as pollinators

**DOI:** 10.1101/2021.04.11.439382

**Authors:** Chen-Chen Zheng, Yi-Bo Luo, Yun-Dong Gao, Peter Bernhardt, Shi-Qi Li, Bo Xu, Xin-Fen Gao

## Abstract

- Flowering plants always attract animals providing rewards or deceptive signals to gain reproductive success. However, there is no well-documented reporting about a pollination mechanism with both rewards and deceptive signals by a same object. We found *Cypripedium wardii* flowers seem to attract visitors by the white pseudopollen-like trichomes on labella in our preliminary field observation.
- To explore the pollination mechanism of *Cypripedium wardii*, especially, the ecological function of the pseudopollen-like trichomes, we conducted field observations, analyses of the traits of visitors and flowers, and breeding system experiments.
- The white trichomes composed by multicellular moniliform hairs on the floral labella played a crucial role to attract pollinators, causing a high natural fruit set ratio in *C. wardii*. We established the direct connection of the white trichomes and real pollen.
- We propose that flowers of *C. wardii* provide pseudopollen to attract suitable bees and hoverflies as pollinators. And our evidence indicate that the pseudopollen owns both deceptive and rewarding ecological functions. Our study provide a clear pollination mechanism with both rewards and deceptive signals by a same object in angiosperm for the first time. However, an inbreeding depression seem to be caused by this strategy. And we speculated that the pollen mimicry strategy with both rewarding and deceptive functions in *C. wardii* may be an adaptation to the habitat fragmentation of this species to gain a reproductive assurance.

## Introduction

Almost all flowering plants, i.e., angiosperm, attract animals providing rewards or deceptive signals to gain reproductive success (Johnson & Schiestl, 2016; Lunau *et al*., 2017; Wester & Lunau, 2017). In addition, some flowers are also reported to adopt both rewards and deceptive signals by different objects, respectively (Meve & Liede, 1994; Bänziger, 1996; Bänziger, 2001; Brodmann *et al*., 2008; Ellis & Johnson, 2010; Gottsberger, 2012; Johnson & Schiestl, 2016; Jiang *et al*., 2020). However, there is no well-documented reporting about a pollination mechanism with both rewards and deceptive signals by a same object. Though without obvious evidence, pseudopollen was the only pollination strategy to be proposed with both reward and deception roles in angiosperm (Davies & Turner, 2004; Davies *et al*., 2013).

Pollen, being an honest signal, is usually used by most flowering plants to reward floral visitors, e.g., bees and hoverflies (Lunau, 2000; Lunau *et al*., 2017). Hence, mainly by color with UV-absorbing visual patterns, pollen- and stamen- mimicries also become common strategies in many angiosperms (Lunau, 2000; Papiorek *et al*., 2016). And even many nectar mimicries are also related to this visual trait (Lunau *et al*., 2020). However, with the existence of accessible pollen, we believe that most pollen- and stamen-mimicries are floral guides actually (Lunau *et al*., 2017) and the pollinators attracted by the dishonest signals can finally gain their initial targets, i.e., true pollen. In fact, deceptive pollination mechanism related to pollen, such as pseudopollen, is uncommon in flowering plant species (Johnson & Schiestl, 2016). But, pseudopollen is frequently proposed in the Orchidaceae family, as the existence of uneatable and uncollectible pollinaria (Sanguinetti *et al*., 2012; Lunau *et al*., 2017; Pansarin & Maciel, 2017).

Pseudopollen, an arenaceous powder, usually occurs on labella of the orchids and is formed by fragmentation of multicellular moniliform trichomes with cells rich in protein and/or starch (Davies & Turner, 2004; Jersáková *et al*., 2006; Davies *et al*., 2013). However, most work about pseudopollen are focusing on its structure, cell contents, and development (Davies *et al*., 2000; Davies *et al*., 2013), lacking fieldwork to figure out its ecology function, i.e., deception and/or reward (Davies *et al*., 2013; Johnson & Schiestl, 2016). To the best of our knowledge, there are only a few orchids with field observations, e.g., *Heterotaxis brasiliensis* (Brieger & Illg) F. Barros, *Maxillaria ochroleuca* G.Lodd. ex Lindl. (Singer & Koehler, 2004), *Mormolyca rufescens* (Lindl.) M.A. Blanco, *Heterotaxis discolo* (G. Lodd. ex Lindl.) Ojeda & Carnevali (Singer *et al*., 2004), *Polystachya flavescens* (Blume) J.J. Sm. (Goss, 1977; Pansarin & Maciel, 2017), *P. estrellensis* Rchb. f., *P. concreta* (Jacq.) Garay & H.R. Sweet (Pansarin & Amaral 2006), *Cyanaeorchis arundinae* (Rchb. f.) Barb. Rodr. (Pansarin & Maciel, 2017), and *Cypripedium subtropicum* S.C. Chen & K.Y. Lang (Jiang *et al*., 2020), which reported the behaviors that bees or hoverflies actively collect/eat the pseudopollen-like trichomes on labella, however, without further experiments to explore the ecology function of the trichomes. As a result, there is a mess about the pseudopollen. Some studies proposed that it’s a deceptive signal to pollinators as the resemblance with real pollen (Davies *et al*., 2000; Jersáková *et al*., 2006; Davies *et al*., 2013). However, some other studies suggest that it’s a reward named edible hairs for the rich contents and the active collection/eating behaviors of visitors (Pansarin & Maciel, 2017; Jiang *et al*., 2020). And some studies suggest that pseudopollen is with both reward and deception pollination function (Davies & Turner, 2004; Davies *et al*., 2013). To solve this mess, we need more detail field and lab work.

Within the genus *Cypripedium*, a model lineage taking deceptive strategies to attract pollinators (Bernhardt & Edens-Meier, 2010), *Cypripedium wardii* Rolfe, an endangered species endemic to the Hengduan Mountains, western China (Chen & Cribb, 2009) was observed with white pseudopollen-like trichomes on floral lip (Fig 1c-e), which were actively visited by bees and hoverflies, in our preliminary field observation in southwest China. The similar trichomes on labella were also reported in *C. subtropicum*, the sisiter species of *C. wardii* (Li et al., 2011), and proposed as rewards to attract hoverfly pollinators (Jiang *et al*., 2020). Hence, we speculate that the trichomes in *C. wardii* may be pseudopollen or edible hairs. Up to date, little is known about the pollination of *C. wardii*.

**Figure 1.**
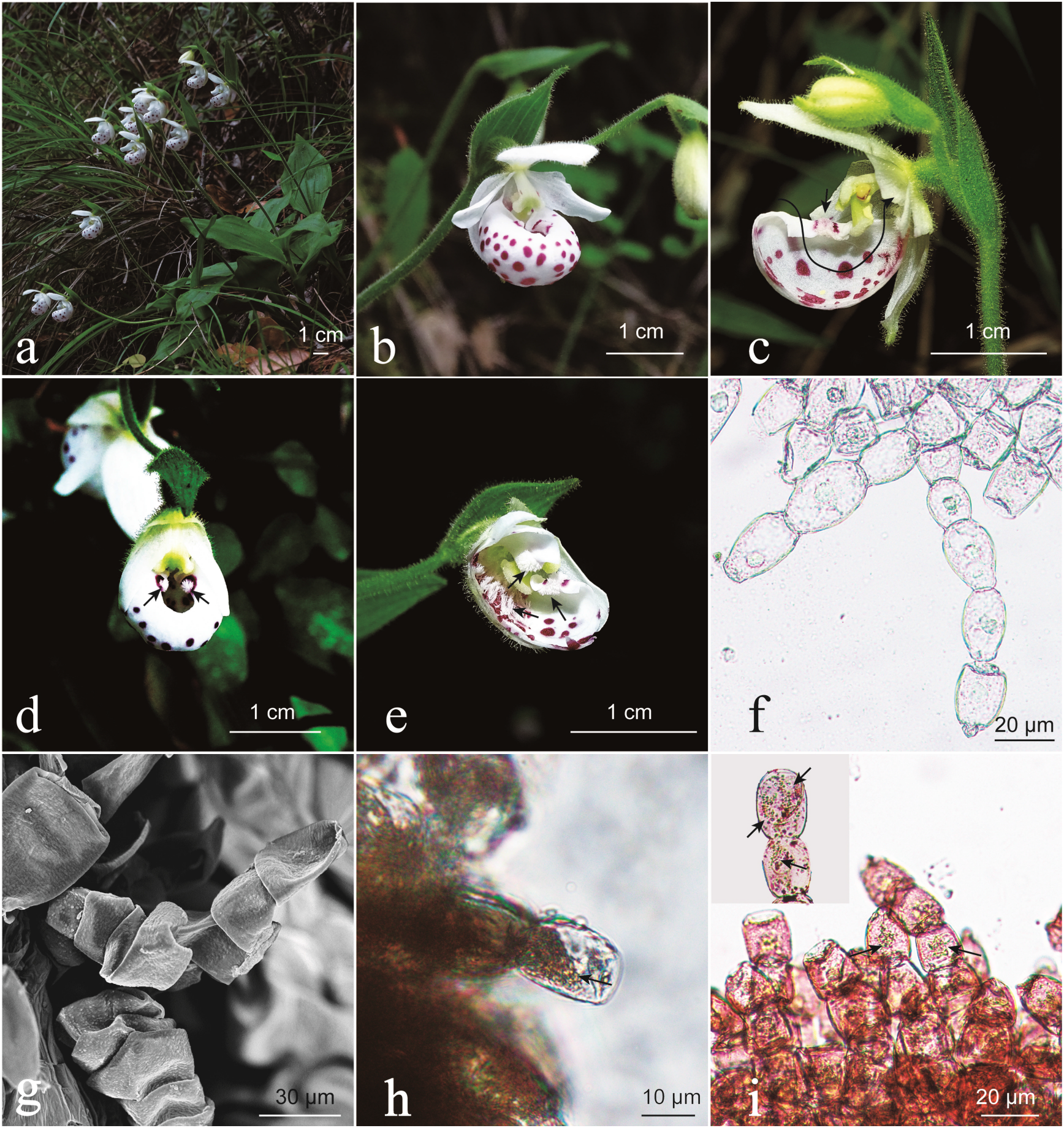
*Cypripedium wardii* and its floral morphology. (a) Natural habitat of *C. wardii*. (b) Front view of a flower. (c) Dissected flower with half the labellum removed. The line indicates the pollination pathway and the white trichomes (arrow) at the beginning of the pollination pathway. (d) The white trichomes on the outer surface of the mouth opening edge (arrow). (e) The white trichomes on the inner surface of the mouth opening edge and inside the labella (arrow). (f) Light microscopy image and (g) scanning electron microscopy (SEM) image of multicellular trichomes on labellum. Light microscopy image of the white hair tufts, showing (h) the contents rich in the composed cell (arrow) and (i) lipids drops in the contents.

In this study, we try to explore the pollination mechanism of *Cypripedium wardii*,especially, to make sure the ecology function, i.e., reward and/or deception, of the pseudopollen-like trichomes. To reach the goal, we conducted two years field observations, analyses of the traits of visitors and flowers (especially for the trichomes on floral labellum), and breeding system experiments. Furthermore, with the result mentioned above, we are trying to discuss the reason *C. wardii* adopt such a pollination mechanism and the trade-off, i.e., what success will gain from the mechanism and the cost will pay for the strategy.

## Material and methods

### Study species and site

*Cypripedium wardii* is a renascent herb and 1-, 2-, or 3-flowered (Fig. 1a) with rate of 7:6:1 (n = 577) in this studied population, correcting the previous description assuming that *C. wardii* is 1- or 2-flowered (Chen & Cribb, 2009). The flowers are small white or creamy white with purple spots on inside labella and around the mouth openings (Fig. 1b-d). And the flowers are with white trichomes on labellum (Fig. 1c-e), an ignored trait in previous description (Chen & Cribb, 2009).

Field observations and wild experiments were conducted in Heba Village, Kangding, Sichuan Province, Southwest China. A large population of *C. wardii* with more than 800 flowering plants was on our site with an area of 15540m^2^. This site was on a limestone mountain with secondary deciduous broad-leaved and coniferous mixed forest at an elevation of 2200–2300 m. The flowers bloom from early of Jun. to mid-July, and the hapaxanthic period was 6.86 ± 3.26 (Mean ± SD, n = 158) days. There were 63 co-flowering species (Table S1) and the dominated co-flowering species were *Berberis wilsoniae* Hemsl., *Calanthe davidii* Franch., *Campylotropis polyantha* (Franch.) Schindl., *Cypripedium lichiangense* S.C. Chen & P.J. Cribb, and *Cotoneaster horizontalis* Decne. The specimens of co-living plants were deposited in Herbarium of Chengdu Institute of Biology, Chinese Academy of Science (CDBI), Chengdu.

### Breeding system experiments

To determine whether the pollinators were necessary for *C. wardii* to produce fruits and seeds and the existence of inbreeding depression, we performed breeding system experiments in 2019 and 2020.

In 2019, following Zheng & Li (2009), we divided marked flowers (n = 333) into three treatment categories before the buds opened: (1) self-pollinated flowers (n = 19), (2) cross-pollinated flowers (n = 23), and (3) naturally pollinated flowers (n = 291). We removed the labella of self-pollinated flowers by a razor blade and hand-pollinated them with the pollinia from their own anthers. We removed the labella of cross-pollinated flowers by a razor blade and hand-pollinated them with the pollinia from flowers located > five meters away. The fruits from the marked plants were counted and collected during mid-October. After backing in lab, we removed and mixed the seeds from every three capsules of different experimental plants, respectively. We identified the types of embryos produced under a stereoscope (Olympus BX43F, Olympus, Japan), such as big, small, aborted, and absent embryos (Jersáková & Johnson, 2006). Moreover, we tested seed viability through pretreatment by soaking in 5% sodium hypochlorite (w/v) for 2 hours and with 1% tetrazolium (w/v) (Van Waes & Debergh, 1986; He, 2010), and then we observed the seeds using a stereoscope (Olympus BX43F, Olympus, Japan): unstained seeds were considered dead seeds and stained seeds (pink and red) were considered viable seeds.

In 2020, we divided marked flowers (n = 395) into four treatment categories before the buds opened: (1) self-pollinated flowers (n = 25), (2) cross-pollinated flowers (n = 27), (3) naturally pollinated flowers (n = 305), and (4) control flowers (n = 38). We bagged the buds of self-pollinated flowers and hand-pollinated them with the pollinia from their own anthers after opening, and then bagged the flowers again and kept them until the end of the flowering periods. We bagged the buds of cross- pollinated flowers and hand-pollinated them with the pollinia from flowers located > five meters away, and then bagged the flowers again and kept them until the end of the flowering periods. We bagged the buds of control flowers and kept them until the end of the flowering periods. The bags were all non-woven tea bags bought online. The fruits of the marked flowers were counted at the end of their flowering periods.

We test the difference of breeding system experiments by χ^2^ test.

### Field observations

Field observations were performed in two flowering seasons, mainly on sunny day, among 9:00–18:00 (daytime in 2019), 10:00–16:00 (daytime in 2020), and 19:00– 24:00 (night in 2020). In total, we observed for 187 hours, 172 h during daytime (112 h in 2019 and 60 h in 2020) and 15 h at night. We recorded approaching, alighting, entering, and escaping behaviours of floral visitors (Nilsson, 1979). Floral visitors were captured using an insect net for identification. The visitor specimens captured were deposited in Herbarium of Chengdu Institute of Biology, Chinese Academy of Science (CDBI), Chengdu. Additionally, we recorded whether the pollinators visited other co-flowering species before/after visiting the flowers of *C. wardii* to determine if *C. wardii* benefited from its co-flowering species.

### Analysis of morphology and contents of the white trichomes

To examine morphological traits, we collected plant material of *Cypripedium wardii* and placed it in 70% alcohol:acetic acid:formaldehyde (8:1:1). After backing in lab, we first observed the floral traits under a light microscope (Olympus BX43F, Olympus, Japan). And then we observed the micro traits under a scanning electron microscope (SEM; Phenom Pro, Netherlands) with dehydration treatments with graded ethanol–isoamyl acetate, then plating the dried materials with gold palladium and observing them at an accelerating voltage of 10 kV (Ren *et al*., 2011).

To make sure whether exist nutrient content in the cells of white trichomes, we conducted histochemical tests for three crucial elements, i.e., protein, lipids, and starch, following Davies *et al*. (2013). We collected fresh flowers of *Cypripedium wardii* from field and keep it at 4 °C environment, then test the three nutrient elements as soon as possible. We removed some trichomes from fresh labellum then placed them on a microscope slide, and dropped a drop of different solutions, then observed the color reaction under a light microscope (Olympus BX43F, Olympus, Japan). For protein, basing on a purple reaction product with 0.2% (w/v) aqueous ninhydrin (indane-1,2,3-trione hydrate) solution, we suggested the existence of protein. For starch, basing on a blue reaction product with iodine–potassium iodide (IKI), we suggested the existence of starch. For lipids, basing on an orange reaction product with a saturated solution of Sudan III in 70% (v/v) ethanol, we suggested the existence of lipids.

### Analysis of attachment on outer surfaces and digestive tracts of pollinators

To make sure whether pollinators collect and/or eat the white trichomes on labella, we examined the outer surfaces and digestive tracts of the main pollinators using SEM with similar treatments mentioned above in *Analysis of shape and content of the white trichomes*.

### Visitors responses to the removal of white trichomes on outer surface of labellum

To test the attraction of the white trichomes, we moved away the visible trichomes being around the outside of mouth openings (i.e., Fig. 1d) of 33 flowers by razor blades, and then observed the visiting behaviors of visitors during daytime, i.e., 10:00–16:00, in 2020. Finally, we evaluated the attraction of the white trichomes basing on the changes of visiting frequency and fruit set ratio between the flowers without outside trichomes and normal flowers in 2020. We test the difference of fruit set ratio by χ^2^ test.

## Results

### Breeding system experiments

Natural fruit set ratio in *Cypripedium wardii* was pretty high, i.e., 82% in 2019 and 76% in 2020, compared with other studied species in genus *Cypripedium* (Bernhardt & Edens-Meier, 2010). All 38 control flowers produced no fruits. Cross-pollinated flowers produced 96% fruits in 2019 and 100% in 2020. Self-pollinated flowers produced 89% fruits in 2019 and 96% in 2020. There were no significant difference between two hand-pollinated flowers (P > 0.05, χ^2^ test). And the seeds among naturally, self- and cross-pollinated flowers in 2019 were main big embryos, i.e., 91%, 79%, and 82%, respectively. However, viability test showed that most seed embryos of all treatments in 2019 were not able to be dyed, i.e., 10% for naturally pollinated flowers, 10% for self-pollinated flowers, and 37% for cross-pollinated flowers. There was no significant difference of the proportions of vigorous seeds (i.e., seeds with embryos dyed red and pink) between naturally and self-pollinated flowers (P > 0.05, χ^2^ test), however, the proportions of vigorous seeds in cross-pollinated flowers was significantly higher than that in both naturally pollinated flowers and self-pollinated flowers (P < 0.001 each, χ^2^ test). The details of results of the breeding system experiments are shown in Table 1.

**Table 1.**
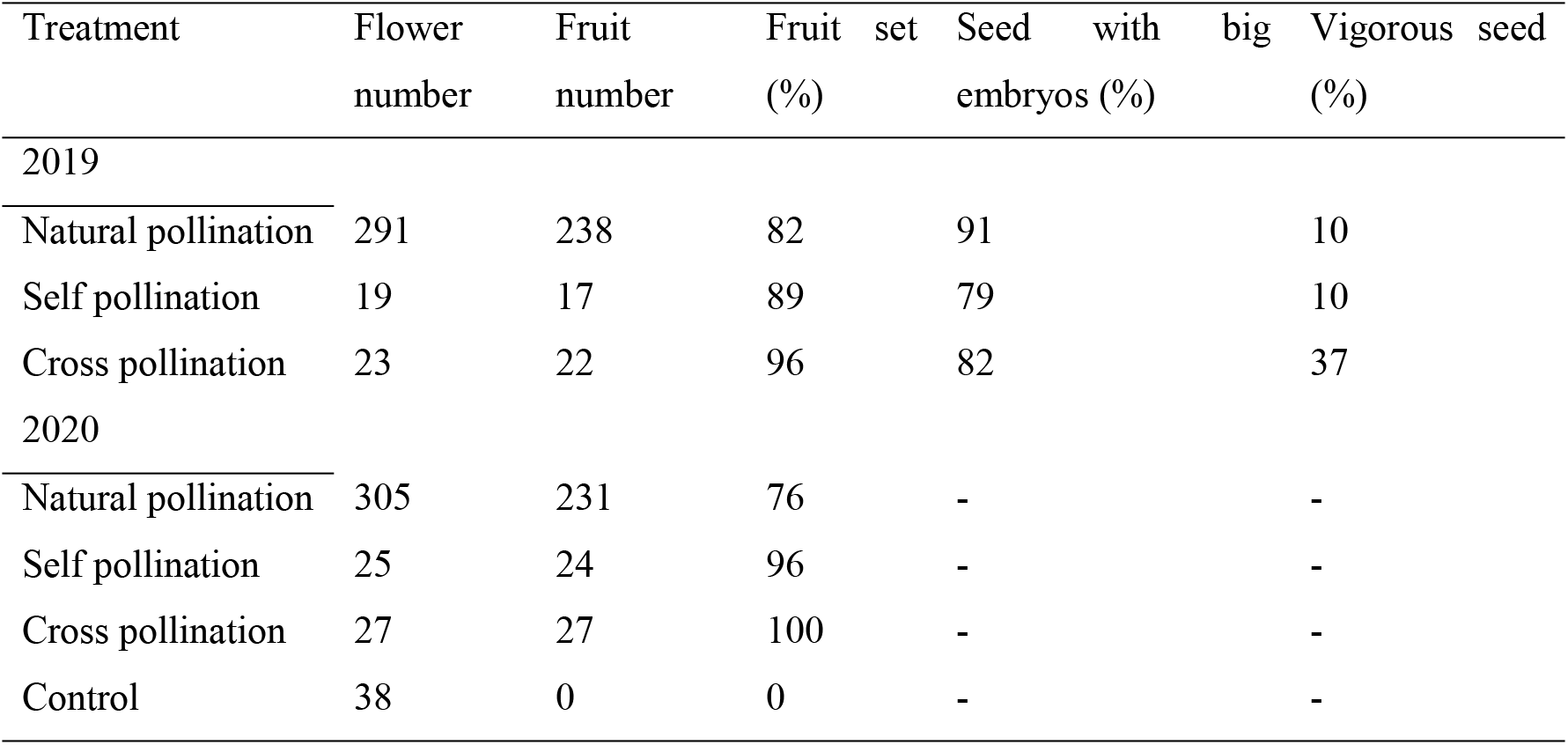
Results of breeding system experiments of *Cypripedium wardii*. Vigorous seeds includes seeds stained pink and red.

### Field observations and pollinators

High visiting frequency of bees and hoverflies to *Cypripedium wardii* flowers were found during daytime, but we did not observe any visitors at night. In total, we observed 277 visits of bees (190) and hoverflies (87) in the whole 172 hours field observation during daytime (details see Fig. S1 and Table S2). The visiting frequecy of visitors was so high that we even observed some spiders (Thomisidae) ambushing on the flowers (e.g., Fig. 2a). Rather than its sister species *C. subtropicum* with a strong floral scent (Jiang *et al*., 2020), there was no obvious human detected scent in *C. wardii*. Basing on whether carrying out the floral pollen, some bees (e.g., Fig. 2b and Video S1) and hoverflies (e.g., Fig. 2e and Video S2) were found to be effective visitors, i.e., pollinators. According to the field observation in 2020, we found 25 pollinators carried out pollen smear and 12 pollintors came to visit flowers with past pollen smear (e.g., Fig. 2c, g). In total, we collected more than 98 specimens of bee and hoverfly visitors, and identified seven species of bees and five species of hoverflies as pollinators of *C. wardii* (Table 2). We did not find any nectar like fluid in the flowers to reward pollinators. However, we found that the pollinators always showed great interest in the white powder-like trichomes distributing on both outside and inside the edge of mouth opening (Fig. 1d-e) and inside the labella (Fig. 1e). Bee pollinators usually flew to the mouth opening directly (e.g., Fig. 1c) and then actively scraped the trichomes both on the edge of mouth opening and inside the labella with their forelegs, and the behaviors similar to biting the white trichomes were also showed during the process of collecting trichomes (e.g., Video S3, S4). The bees seemed to be guided into the labella by the trichomes on the inside the edge of mouth opening (e.g., Fig. 1c-e). Hoverfly pollinators also flew to the mouth opening directly (e.g., Fig. 2g) and then showed the behaviors of proboscis extension to the trichomes on the edge of mouth opening (e.g., Video S5-S7 and Fig. 2d), and some were guided into the labella by the inner trichomes finally (e.g., Video S8). We never found dead bees in the labella. However, dead hoverflies were found in some labella, which might be related with their oversized bodies. Most pollinators visited the flowers following the planned pollination pathway (Fig. 1c), however, we also observed that some bee pollinators escaped the pouch from mouth opening and entered the pouch from one of the back holes. Some pollinators, including both bees and hoverflies, were observed to re-visit the same flower and/or visit another flower of *C. wardii* after escaping from one flower (e.g., Video S1, S5). In total, we observed 16 continuous visits in 2020. In addition, both bee and hoverfly pollinators also visited other co-flowering plants after/before visiting *C. wardii*, such as *Gentiana rubicunda* Franch. (Fig. 2i-j), *Pedicularis* sp. (Fig. 2k), and *Leptodermis potanini* Batalin (Fig. 2m), indicating a benefit from co-flowering plants might help the pollination of *C. wardii*. We never found the egg-laying behaviors of pollinators and eggs on the flowers during the whole field observational period.

**Figure 2.**
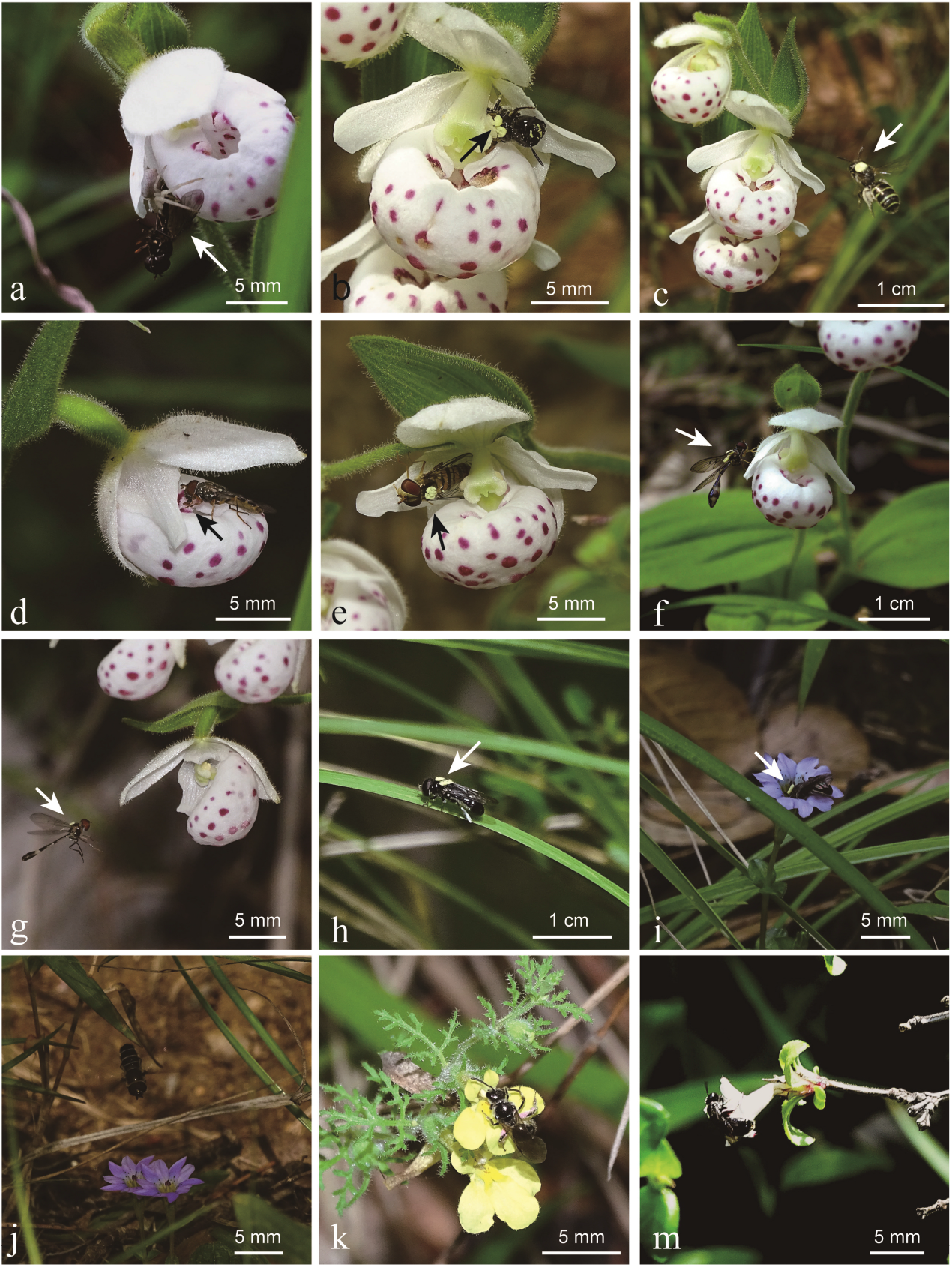
Field records of *Cypripedium wardii* visitors. (a) A spider (Thomisidae) ambushing on the flowers (arrow). (b) A bee pollinator (*Ceratina japonica*) carrying out pollen smear (see arrow) from one of the back exits. (c) A bee pollinator (*Ceratina japonica*) with past pollen smear (see arrow) visiting a *C. wardii* flower. (d) A hoverfly behaving the proboscis extension to the trichomes on the edge of mouth opening (arrow). (e) A hoverfly pollinator (*Episyrphus balteata*) carrying out pollen smear (see arrow) from one of the back exits. (f) A hoverfly pollinator (*Baccha maculata*) carried out pollen smear (see arrow) and rested on the labellum. (g) A hoverfly pollinator (*Baccha maculata*) with past pollen smear (see arrow) visiting a *C. wardii* flower. (h) A hoverfly pollinator (*Eumerus lucidus*) with pollen smear (see arrow) resting on a leaf. (i) A pollinator with pollen smear (see arrow) visiting a *Gentiana rubicunda* flower after visiting a *C. wardii* flower. (j) A hoverfly visitor visiting a *Gentiana rubicunda* flower. (k) A bee visitor visiting a *Pedicularis* sp. flower. (m) A bee visitor visiting a *Leptodermis potanini* flower.

**Table 2.** Pollinators of *Cypripedium wardii*.

| Species | Family |
| --- | --- |
| Bees |  |
| *Ceratina chinensis* Wu | Apidae |
| *C. japonica* Cockerell | Apidae |
| *Lasioglossum compressum* Blüthgen | Apidae |
| *L. macrurum* Cockerell | Apidae |
| *L. proximatum* Smith | Apidae |
| *L. sinicum* Blüthgen | Apidae |
| *Seladonia varentzowi* Morawitz | Apidae |
| Hoverflies |  |
| *Baccha maculata* Walker | Syrphidae |
| *Episyrphus balteata* De Geer | Syrphidae |
| *Eumerus lucidus* Loew | Syrphidae |
| *Melanostoma orientale* Wiedemann | Syrphidae |
| *Sphaerophoria rueppelli* Wiedemann | Syrphidae |

### Morphology and contents of the white trichomes

SEM and light microscopy images showed that the white trichomes on the floral labella (Fig. 1d-e) were formed by many multicellular moniliform hairs (Fig. 1f-g), corresponding with the general description of pseudopollen (Davies & Turner, 2004; Jersáková *et al*., 2006; Davies *et al*., 2013). In addition, light microscopy images showed that formed cells of the trichomes were rich in some unknown contents (Fig. 1h). And histochemical tests showed that lipids (Fig. 1i) were rich in the contents of trichomes cells. However, our results did not show the existence of protein and starch, which were usually reported in the pseudopollen of other orchid (Davies & Turner, 2004; Davies *et al*., 2013).

### Attachment on outer surfaces and digestive tracts of pollinators

In total, we examined 12 bee and five hovefly specimens. The pollen smear attached to the pollinators (e.g., Fig. 2b-c, e-i), i.e., pollen of *Cypripedium wardii*, was shown in Fig. 3a. We found that all kinds of pollen mixed with the formed cells of the white trichomes (COT) attached to the hind legs of all detected bee specimens (e.g., Fig. 3b). Though the pollen types on different bee individuals were diverse (e.g., Fig. 3c), the existence of COT was consistent (Table S3). In total, 15 kinds of pollen grains were found on the hind legs of bee pollinators (Fig. S2). In addition, we also observed COT on the mouthparts (e.g., Fig. 3d) and in the guts (e.g., Fig. 3e) of some bee specimens. The identification of COT on detected specimens was based on the shape and ornamentation, compared with the shape and ornamentation of the white trichomes on labellum (Fig. 1g). We never observed COT attached to the outer surfaces of the examined hoveflies, however, COT mixed with all kinds of pollen were found in the guts of all five detected specimens (e.g., Fig. 3f). The SEM images of outer surfaces and digestive tracts of the examined pollinator specimens indicated that all the pollinators collected/ate the white trichomes as food, corresponding to our field observations (e.g., Fig. 1d and Video S3-S8). In addition, the phenomenon that COT mixed with all kinds of pollen implied that bee and hoverfly pollinators treated the white trichomes as pollen.

**Figure 3.**
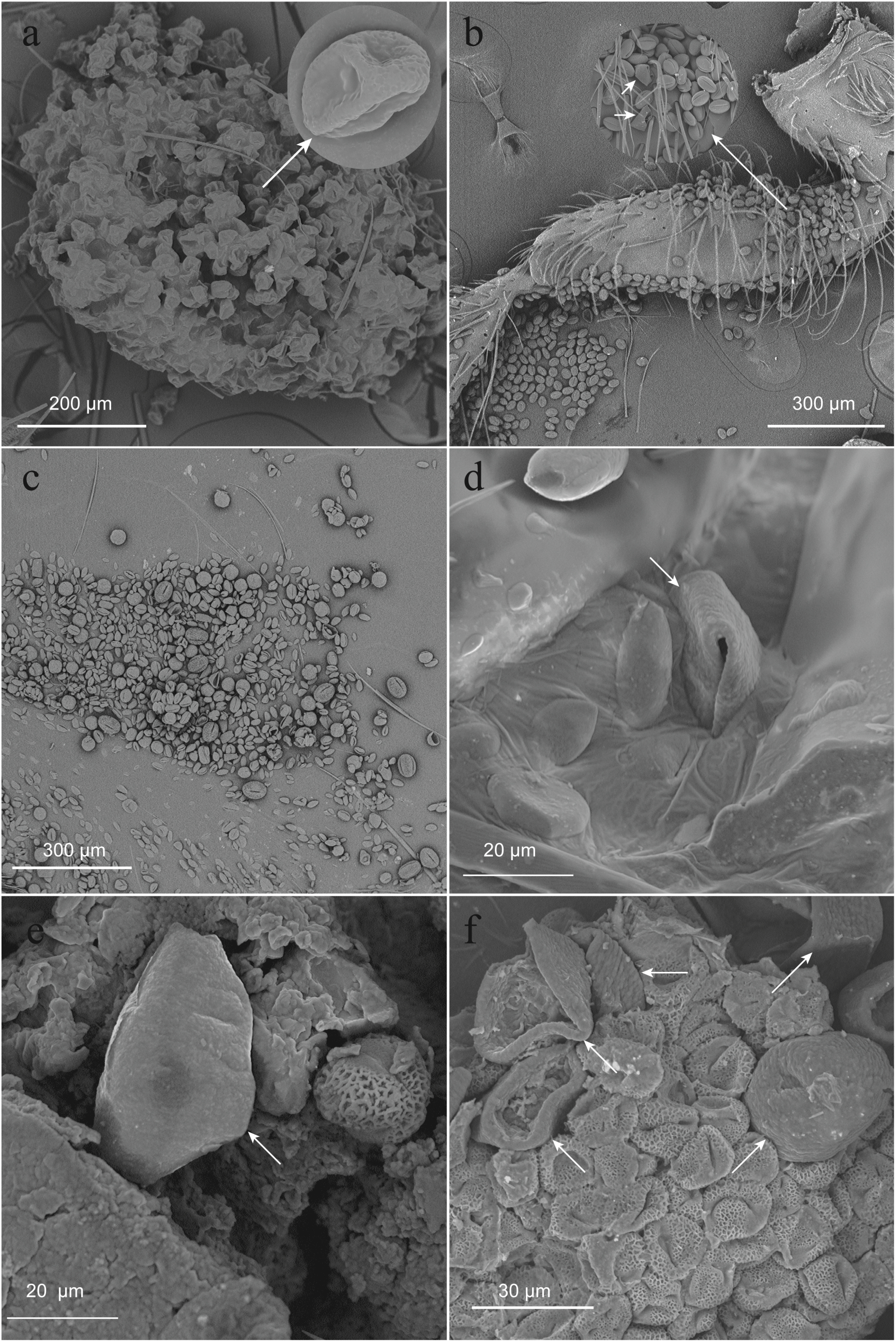
Scanning electron microscopy (SEM) images of bee and hoverfly pollinators. (a) The pollen smear on a pollinator and the morphology of a single pollen grain (see arrow). (b) The formed cells of the white trichomes (COT, see arrow) mixed with pollen attached to a hind leg of bee pollinator. (c) The diverse pollen mixed with COT attached to a hind leg of bee pollinator. (d) A formed cell of the white trichomes (see arrow) attached to the mouth part of a bee pollinator. (e) A formed cell of the white trichomes (see arrow) in the gut of a bee pollinator. (f) COT (see arrow) mixed with pollen in the gut of a hoverfly pollinator.

### Visitors responses to the removal of white trichomes on outer surface of labellum

In total, we observed for 30 hours to test the behaviour responses of visitors to the removal of white trichomes on outer surface of labellum (i.e., Fig. 1d). We did not observe an obvious change of visiting behaviors of bees to the flowers with removal of white trichomes on outer surface of labellum (Table S4), comparing with that in the normal flowers (Table S2). Bee visitors would fly to the mouth opening and enter the labella directly, and then actively collected the trichomes on the inner surface of edge of mouth opening and inside the labella (e.g., Video S9, S10). And we also observed eight continuous bee visits (e.g., Video S10), i.e., re-visiting the same flower and/or visiting another flower of *C. wardii* after escaping from one flower. In total, we observed 32 bee visitors entering labella. There were not an obvious difference between bee visiting frequencies to flower with (1.63/h) and without (1.78/h) the removal of white trichomes on outer surface of labella in 2020. We speculated that the reasons why there were no changes of bee visiting behaviors were that the experienced bees had learnt the existence of the white trichomes in the labella by previous experiments and the bees were easy to escape from the trap labella. However, we never observed that hoverfly visitors entered the labella with removal of white trichomes on outer surface of labella, though there were ten visiting records (Table S4). It seemed that the removal of white trichomes on outer surface of labellum only influenced the visits of hoverflies, as the hoverflies needed the outer trichomes to guide into the labella (e.g., Video S8). However, we only got 15 fruits in the 33 examined flowers. There was a significant reduction of the fruit set ratio in examined flowers (45%) comparing with that in the nature pollinated (76%) in 2020 (P < 0.001, χ^2^ test). In brief, the white trichomes on the labella of *Cypripedium wardii* were necessary to attract its visitors.

## Discussion

None of control flowers produced fruits, indicating no autogamous self-pollination in *Cypripedium wardii* and pollinators were necessary for its sexual propagation. However, unlike most of the other reported species in *Cypripedium*, e.g., *C. fargesii*(Ren *et al*., 2011), *C. wardii* is not limited by its pollinators to produce fruits in the wild. Hence, the pollination strategy adopted by *C. wardii* was successful to attract its pollinators.

Undoubtedly, the white trichomes composed by multicellular moniliform hairs on the floral labella (Fig. 1d-g) played a crucial role to attract pollinators, basing on our field and lab work. In this study, we propose a rare pollination strategy combined with both reward and deception (Johnson & Schiestl, 2016; Jiang *et al*., 2020) in *Cypripedium*, a genus being regard as model lineage of unrewarding orchid flowers (Bernhardt & Edens-Meier, 2010). Flowers of *C. wardii* provide pseudopollen (i.e., the white trichomes shown in Fig. 1d-g), a general food-deceptive strategy (Jersáková *et al*., 2006; Johnson & Schiestl, 2016), to attract suitable bees (e.g., Fig. 2b-c) and hoverflies (e.g., Fig. 2e-h) as pollinators. For the deceptive ecology function of pseudopollen, previous studies mainly base on visual evidence in other studied orchids, i.e., resembling pollen in appearance (e.g., Davies *et al*., 2000; Davies & Turner, 2004; Singer *et al*., 2004; Singer & Koehler, 2004; Davies *et al*., 2013). In a few studies, the researchers also observed pseudopollen-collecting behaviors of bee visitors and treated them as cheating evidence (e.g., Singer *et al*., 2004; Singer & Koehler, 2004; Pansarin & Amaral 2006; Pansarin & Maciel, 2017). However, all of these previous work never established a direct connection of pseudopollen and real pollen. Here, we provide more direct evidence in *C. wardii* that flowers attract generalized pollen collectors, i.e., bees and hoverflies (e.g., Fig. 2i-m) to scraped/ate the pseudopollen on labella (e.g., Fig. 2d; Video S3-8) with supporting of lab work. Our SEM images of pollinators specimens showed that pseudopollen mixed with all kinds of pollen was on the hind legs of all detected bee pollinaters (e.g., Fig. 3b and S2; Table S3) and was rich in the guts of all detected hoverfly pollinators and part bee pollinators (e.g., Fig. 3e-f), establishing a direct connection of pseudopollen and real pollen. For the rewarding ecology function of pseudopollen, also called edible hairs (Pansarin & Maciel, 2017), previous studies were mainly based on the nutrient contents (e.g., protein bodies, lipid droplets, and starch grains) in the composed hair cells for studied orchids (e.g., a review in Pansarin & Maciel, 2017; Jiang *et al*., 2020). In *C. subtropicum*, the hoverfly pollinators’ behaviors of proboscis extension to the similar white trichomes on labella were observed, which were treated as important evidence for rewarding ecology function of pseudopollen (Jiang *et al*., 2020). In *C. wardii*, we also found the nutrient trait of pseudopollen (Fig. 1h-i). In addition, we clearly proved the collecting/eating pseudopollen habits of bee and hoverfly pollinators with both field and lab work (e.g., Fig., 2d, 3b-f, and S2; Video S3-8; Table S3).

With pseudopollen, we first discovered a clear pollination strategy attracting both flies and bees as pollinators in the genus *Cypripedium*, which are usually reported to be pollinated by specific bees or flies (Bernhardt & Edens-Meier, 2010; Edens-Meier *et al*., 2014). In addition, our study might provide an example to help with solving the mess of ecology function of pseudopollen/edible hairs (Davies *et al*., 2000; Davies & Turner, 2004; Jersáková *et al*., 2006; Davies *et al*., 2013; Pansarin & Maciel, 2017; Jiang *et al*., 2020). In this study, we propose that *C. wardii* provide a same object, i.e., pseudopollen, to cheat and reward pollinators. To the best of our knowledge, this is the first reporting about a pollination mechanism with both rewarding and deceptive function by a same object in angiosperm. Davies and Turner (2004) also proposed a similar mechanism for *Dendrobium unicum* Seidenf. basing on the discovery of nutrient contents of the pseudopollen. However, they provided no further evidence, such as pollination ecology evidence.

*Cypripedium wardii* seem to get an obvious reproductive success with the pollination strategy of pseudopollen, basing on the high conversion ratio of flowers into fruits (78%-82%; Table 1) and high ratio of seeds with big embryos (91%; Table 1). However, there also seem to be an inbreeding depression in *C. wardii* as only 10% seeds in naturally pollinated fruits are vigorous, showing no significant difference with self-pollinated fruits (P > 0.05, χ^2^ test; Table 1). And the ratio of seeds with viability in cross-pollinated fruits are significantly higher than that in self-pollinated and naturally pollinated fruits, respectively (P < 0.001 each, χ^2^ test; Table 1). Our field observations also showed that both bee and hoverfly pollinators would re-visit the same flower and/or visit another flower of *C. wardii* after escaping from one flower (e.g., Video S1, S5), making inbreeding possible in nature. Inbreeding caused by pollinators as mediums is an uncommon phenomenon in *Cypripedium* (Edens-Meier *et al*., 2014) and it is generally assumed that one of the main goal of specialized floral mimicry is avoiding inbreeding depression (Johnson & Schiestl, 2016). However, why *C. wardii* has evolved such a pollination strategy? We think it may be related with habitat fragmentation of *C. wardii*, as we never found another population of *C. wardii* near our study site and the nearest population was found on another mountain, more than ten kilometers away from our study site (unpublished data). With both reward and deception ecology functions, pseudopollen in *C. wardii* may be an adaptation to habitat fragmentation to gain a reproductive assurance, which usually is accomplished by autogamous self-pollination for adapting the lacking of pollinators (Wang *et al*., 2004; Liu *et al*., 2006; Fan *et al*., 2012; Johnson & Schiestl, 2016).

In this study, we conducted some preliminary work about the pollination ecology of *Cypripedium wardii* and gained our present conclusion. However, there are also some lacks and important contents are worth further study. Firstly, we suggest that the pseudopollen in *C. wardii* mimics pollen mainly by the visual signal, i.e., the ultraviolet light (UV) absorbing visual patterns (Lunau, 2000; Papiorek *et al*., 2016), but we failed to confirm this as the absence of an UV-light camera. We encourage other researchers to confirm this hypothesis, getting a similar results as shown in Fig 1B by Pansarin & Maciel (2017). Secondly, we did not conduct a scent analysis for *C. wardii* flowers as little is known about the general function of pollen scent in attracting visitors (Lunau, 2000; Lunau *et al*., 2017; Wester & Lunau, 2017), which is significative to explore the chemical information related to the interaction of pollen (or pollen mimicry, e.g., pseudopollen) and animal pollinators. Thirdly, we test the three main nutrient contents, i.e., starch, protein, and lipids, for the contents of pseudopollen and only found the existence of lipids (e.g., Fig. 1i). However, we cannot draw a conclusion that the absence of protein as there is no satisfactory histochemical test for protein (Davies *et al*., 2013). Fourthly, inbreeding depression in *C. wardii* was just based on the seed viability test. But, it is not enough and we need more evidence to convince it, such as seed germination experiments. Fifthly, we found the habitat fragmentation trait in our study site, basing on the distribution of *C. wardii*. However, we know little about the selected environment factors, which are important to protect this endangered species and need to be researched urgently.

In conclusion, we propose a pollination mechanism (i.e., pseudopollen) combined both reward and deception in *C. wardii* to attract suitable generalized pollen collectors (i.e., bees and hoverflies) as pollinators. Pseudopollen strategy in *C. wardii* is successful to attract pollinators, causing a pretty high fruit set ratio in nature. However, there are an inbreeding depression caused by this pollination strategy, i.e., most seeds produced in nature are not viable. The pollen mimicry strategy with both rewarding and deceptive function in *C. wardii* may be an adaptation to the habitat fragmentation of this species to gain a reproductive assurance.

## Acknowledgements

We thank Dr. H.L. Xu for identification of insects. We thank Dr. L.B. Zhang, Dr. Z.X. Ren, Dr. X.H. Xiong, and J.C. Li for providing help in this manuscript. This work was supported by the Science and Technology Basic Work (Grant No. 2017FY100104)

## Supplementary materials

### Supplementary tables

**Table S1.**
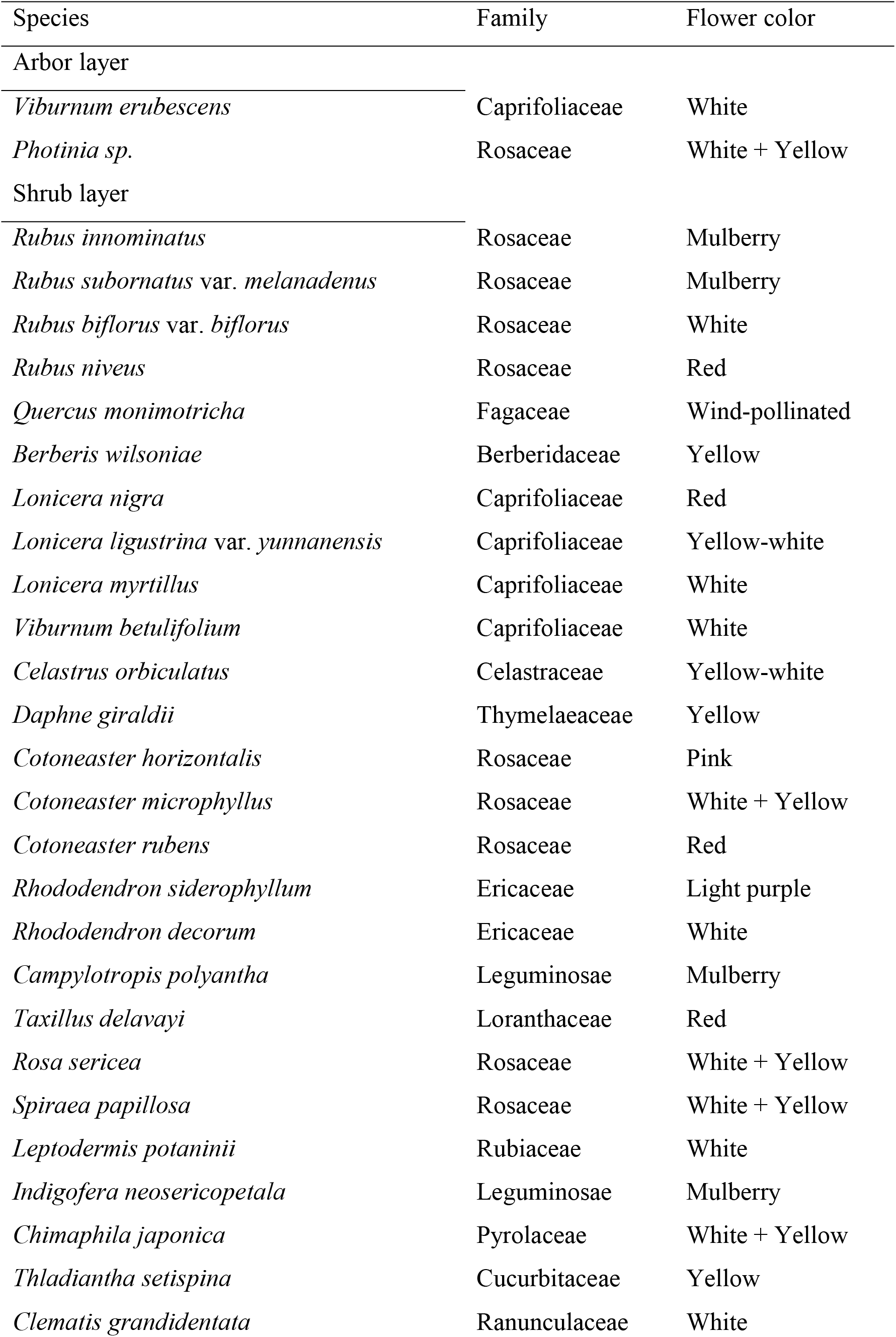
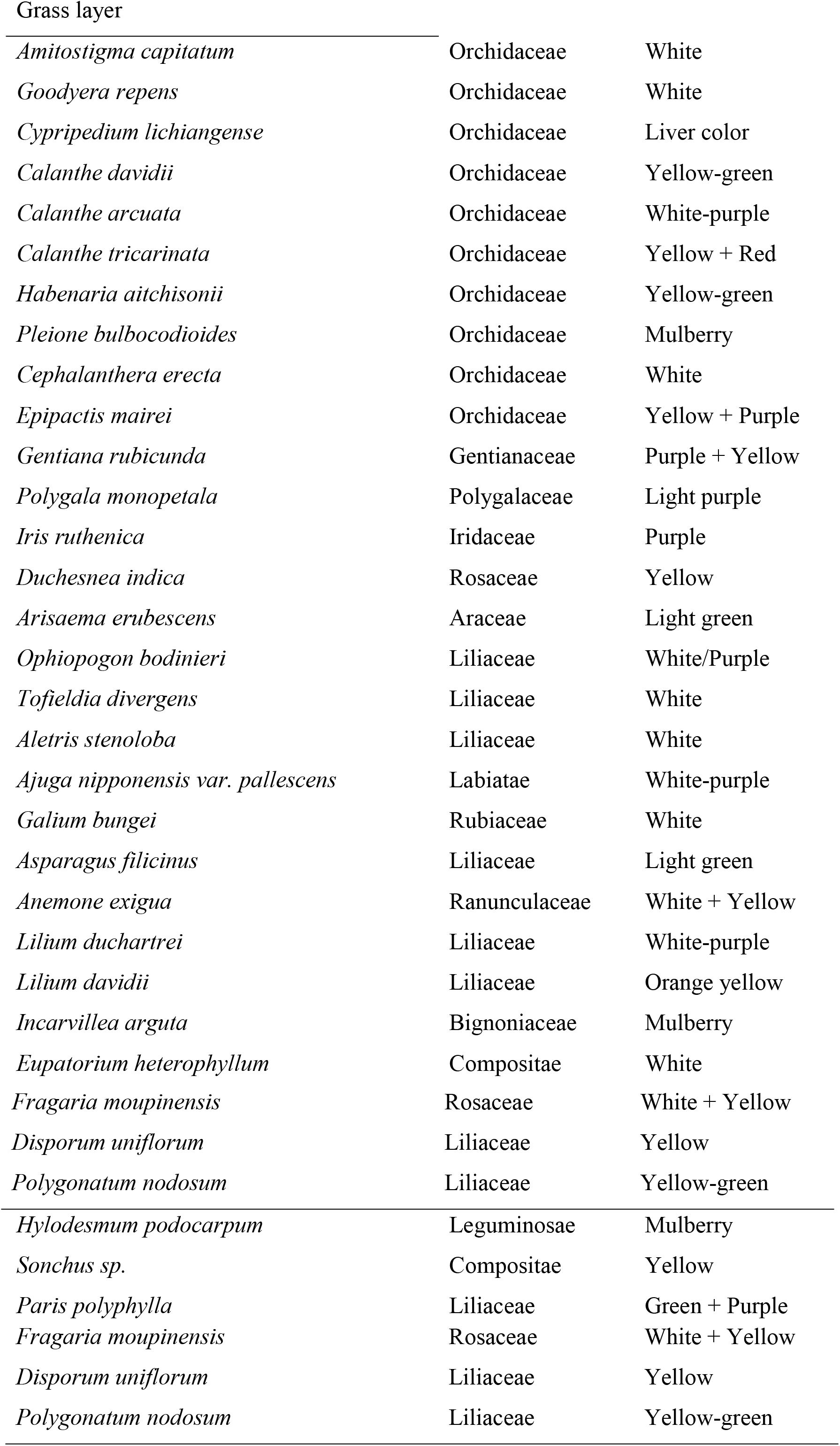
Co-flowering species with *Cypripedium wardii*. the color front of “+” means peripheral color of flowers, the color back of “+” means inside color of flowers.

| Species | Family | Flower color |
| --- | --- | --- |
| Arbor layer |  |  |
| *Viburnum erubescens* | Caprifoliaceae | White |
| *Photinia* sp. | Rosaceae | White + Yellow |
| Shrub layer |  |  |
| *Rubus innominatus* | Rosaceae | Mulberry |
| *Rubus subornatus* var. *melanadenus* | Rosaceae | Mulberry |
| *Rubus biflorus* var. *biflorus* | Rosaceae | White |
| *Rubus niveus* | Rosaceae | Red |
| *Quercus monimotricha* | Fagaceae | Wind-pollinated |
| *Berberis wilsoniae* | Berberidaceae | Yellow |
| *Lonicera nigra* | Caprifoliaceae | Red |
| *Lonicera ligustrina* var. *yunnanensis* | Caprifoliaceae | Yellow-white |
| *Lonicera myrtillus* | Caprifoliaceae | White |
| *Viburnum betulifolium* | Caprifoliaceae | White |
| *Celastrus orbiculatus* | Celastraceae | Yellow-white |
| *Daphne giraldii* | Thymelaeaceae | Yellow |
| *Cotoneaster horizontalis* | Rosaceae | Pink |
| *Cotoneaster microphyllus* | Rosaceae | White + Yellow |
| *Cotoneaster rubens* | Rosaceae | Red |
| *Rhododendron siderophyllum* | Ericaceae | Light purple |
| *Rhododendron decorum* | Ericaceae | White |
| *Campylotropis polyantha* | Leguminosae | Mulberry |
| *Taxillus delavayi* | Loranthaceae | Red |
| *Rosa sericea* | Rosaceae | White + Yellow |
| *Spiraea papillosa* | Rosaceae | White + Yellow |
| *Leptodermis potaninii* | Rubiaceae | White |
| *Indigofera neosericipetala* | Leguminosae | Mulberry |
| *Chimaphila japonica* | Pyrolaceae | White + Yellow |
| *Thladiantha setispina* | Cucurbitaceae | Yellow |
| *Clematis grandidentata* | Ranunculaceae | White |
| Grass layer |  |  |
| *Amitostigma capitatum* | Orchidaceae | White |
| *Goodyera repens* | Orchidaceae | White |
| *Cypripedium lichiangense* | Orchidaceae | Liver color |
| *Calanthe davidii* | Orchidaceae | Yellow-green |
| *Calanthe arcuata* | Orchidaceae | White-purple |
| *Calanthe tricarinata* | Orchidaceae | Yellow + Red |
| *Habenaria aitchisonii* | Orchidaceae | Yellow-green |
| *Pleione bulbocodioides* | Orchidaceae | Mulberry |
| *Cephalanthera erecta* | Orchidaceae | White |
| *Epipactis mairei* | Orchidaceae | Yellow + Purple |
| *Gentiana rubicunda* | Gentianaceae | Purple + Yellow |
| *Polygala monopetala* | Polygalaceae | Light purple |
| *Iris ruthenica* | Iridaceae | Purple |
| *Duchesnea indica* | Rosaceae | Yellow |
| *Arisaema erubescens* | Araceae | Light green |
| *Ophiopogon bodinieri* | Liliaceae | White/Purple |
| *Tofieldia divergens* | Liliaceae | White |
| *Aletris stenoloba* | Liliaceae | White |
| *Ajuga nipponensis* var. *pallescens* | Labiatae | White-purple |
| *Galium bungei* | Rubiaceae | White |
| *Asparagus filicinus* | Liliaceae | Light green |
| *Anemone exigua* | Ranunculaceae | White + Yellow |
| *Lilium duchartrei* | Liliaceae | White-purple |
| *Lilium davidii* | Liliaceae | Orange yellow |
| *Incarvillea arguta* | Bignoniaceae | Mulberry |
| *Eupatorium heterophyllum* | Compositae | White |
| *Youngia* sp. | Compositae | Yellow |
| *Polygonum viviparum* | Polygonaceae | White |
| *Cardamine* sp. | Cruciferae | Mulberry |
| *Hylodesmum podocarpum* | Leguminosae | Mulberry |
| *Sonchus* sp. | Compositae | Yellow |
| *Paris polyphylla* | Liliaceae | Green + Purple |
| *Fragaria moupinensis* | Rosaceae | White + Yellow |
| *Disporum uniflorum* | Liliaceae | Yellow |
| *Polygonatum nodosum* | Liliaceae | Yellow-green |

**Table S2.** The visiting types of bees and hoverflies in the whole 172 hours field observation during daytime. “Only approaching” means that visitors approach flowers but never land on them; “Only landing” means that visitors land on flowers but never enter the labella; “Only entering” means that visitors enter the labella but never carried out pollen smear; “Carrying out pollen smear” means that visitors enter the labella and carry out pollen smear.

| Types | Only approaching | Only landing | Only entering | Carrying out pollen smear |
| --- | --- | --- | --- | --- |
| bees | 4 | 46 | 110 | 32 |
| hoverflies | 7 | 58 | 19 | 1 |

**Table S3.** The attachments on the hind legs of examined bee pollinators. “COA” represents the formed cells of the white trichomes. The letter “Y” means the existence of COA. The letter “N” means the absence of COA.

| Specimens | Pollen types | Existence of COA |
| --- | --- | --- |
| 1 (*Lasioglossum sinicum*) | 6 | Y |
| 2 (*Lasioglossum sinicum*) | 4 | Y |
| 3 (*Lasioglossum sinicum*) | 4 | Y |
| 4 (*Ceratina chinensis*) | 4 | Y |
| 5 (*Ceratina japonica*) | 5 | Y |
| 6 (*Ceratina japonica*) | 2 | Y |
| 7 (*Ceratina japonica*) | 1 | Y |
| 8 (*Ceratina japonica*) | 2 | Y |
| 9 (*Lasioglossum sinicum*) | 3 | Y |
| 10 (*Lasioglossum sinicum*) | 3 | Y |
| 11 (*Lasioglossum compressum*) | 2 | Y |
| 12 (*Seladonia varentzowi*) | 3 | Y |

**Table S4.** The visiting types of bees and hoverflies to flowers with removal of white trichomes on outer surface of labellum in the whole 30 hours field observation during daytime. “Only approaching” means that visitors approach flowers but never land on them; “Only landing” means that visitors land on flowers but never enter the labella; “Only entering” means that visitors enter the labella but never carried out pollen smear; “Carrying out pollen smear” means that visitors enter the labella and carry out pollen smear.

| Types | Only<br>approaching | Only<br>landing | Only<br>entering | Carrying out pollen<br>smear |
| --- | --- | --- | --- | --- |
| bees | 1 | 16 | 22 | 10 |
| hoverflies | 2 | 8 | 0 | 0 |

### Supplementary figures

**Figure S1.**
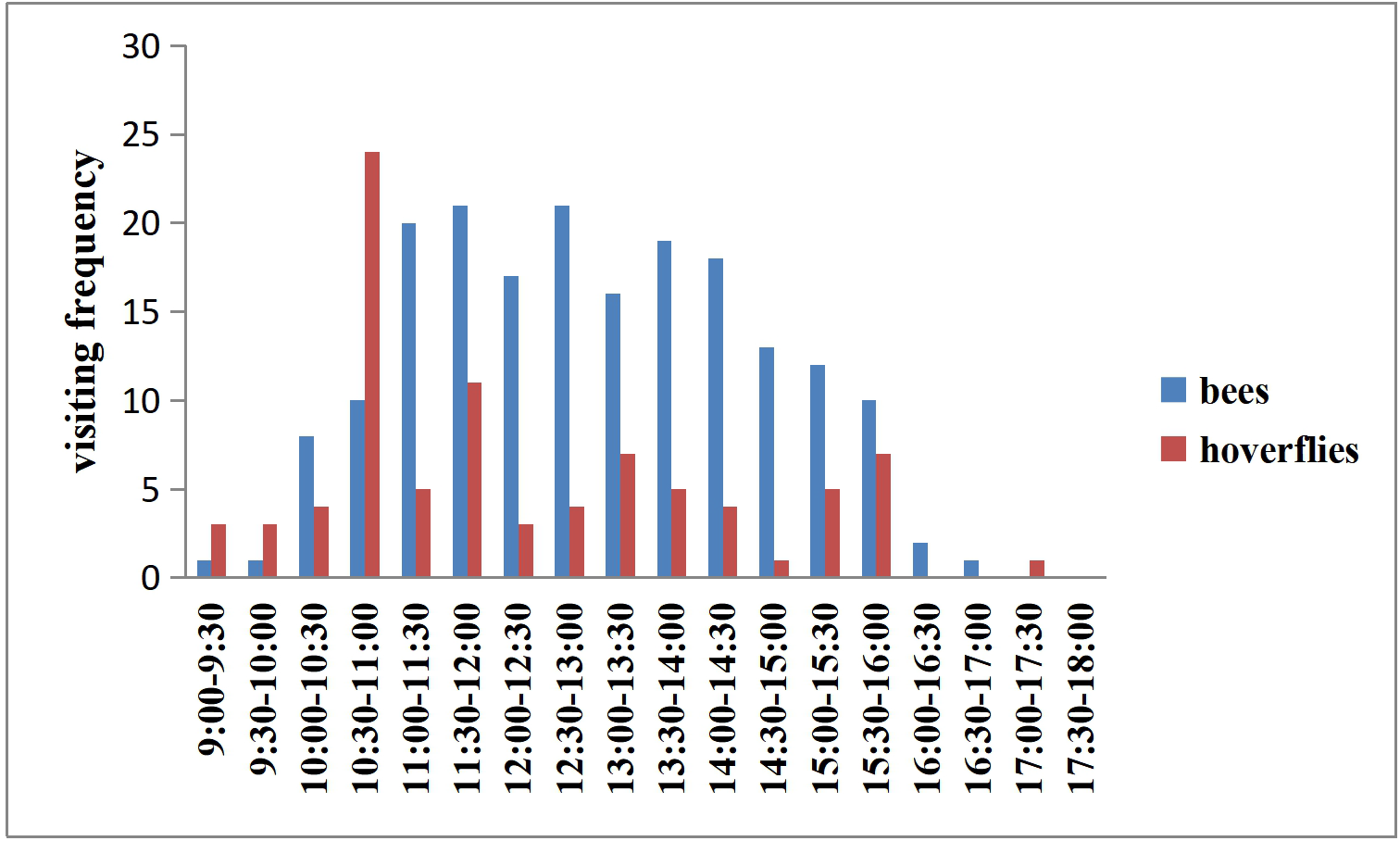
The visiting records of bees and hoverflies in the whole 172 hours field observation during daytime.

**Figure S2.**
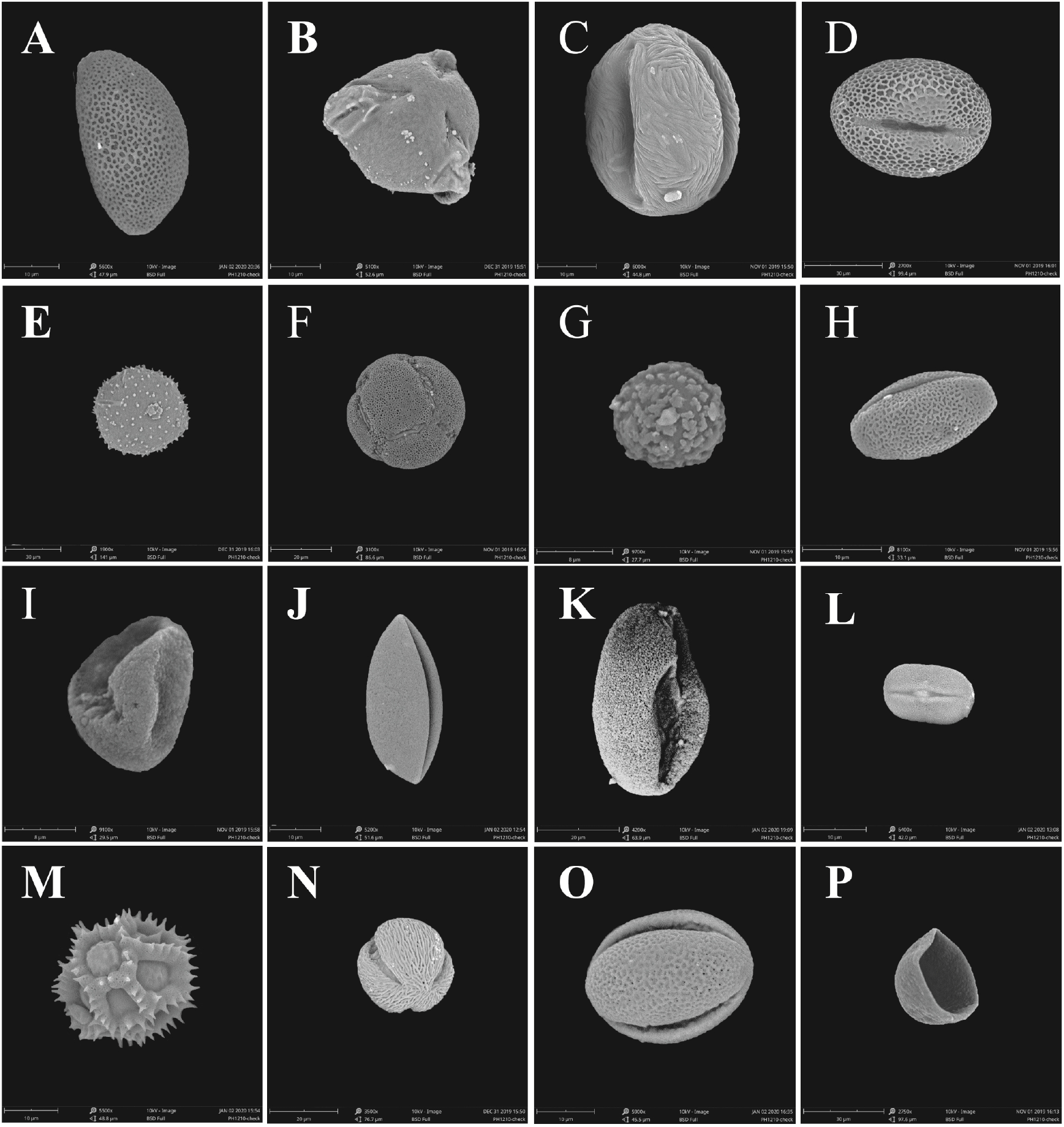
Scanning electron microscopy (SEM) images of the pollen types and the formed cell of the white trichomes attached to the hind legs of the bees. A-O. 15 kinds of different pollen. P. the formed cell of the white trichomes attached to the hind legs of the bees.

### Supplementary videos

Video S1-S10 Visiting records of bee and hoverfly visitors.

## References

Bänziger, H. 1996. Pollination of a flowering oddity: Rhizanthes zippelii (Blume) Spach (Rafflesiaceae). Natural History Bulletin of the Siam Society, 44, 113–142.

Bänziger, H. 2001. Studies on the superlative deceiver: Rhizanthes Dumortier (Rafflesiaceae). Bulletin of the British Ecological Society, 32, 36–39.

Bernhardt P., Edens-Meier R. 2010. What we think we know vs. what we need to know about orchid pollination and conservation: *Cypripedium* L. as a model lineage. Botanical Review, 76(2), 204–219.

Brodmann J., Twele R., Francke W., HoLzler G., Zhang Q.H., Ayasse M. 2008. Orchids mimic green-leaf volatiles to attract prey-hunting wasps for pollination. Current Biology, 18(10), 740–744.

Chen X.Q., Cribb P.J. 2009. Cypripedium L. In Wu ZY, Raven PH, Hong DY, eds. Flora of China. Vol. 25. Beijing: Science Press et St. Louis: Missouri Botanical Garden Press.

Davies K.L., Turner M.P. 2004. Preudo pollen in *Dendrobium unicum* Seidenf. (Orchidaceae): reward or deception? Annals of Botany, 94(1), 129–132.

Davies K.L., Malgorzata S., Magdalena K. 2013. Dual deceit in preudopollen producing *Maxillaria* s.s. (Orchidaceae: Maxillariinae). Botanical Journal of the Linnean Society, 173(4), 744–763.

Davies K.L., Winters C., Turner M.P. 2000. Pseudopollen: its structure and development in maxillaria(orchidaceae). Annals of Botany, 88, 887–895.

Edens-Meier R., Yi-bo L., Pemberton R., Bernhardt P. (2014) Pollination and floral evolution of slipper orchids (Subfamily Cypripedioideae). In: Edens-Meier, R. and Bernhardt P. (editors) eds. Darwin’s Orchids: Then and Now. University of Chicago Press, USA, 265–290.

Ellis A.G., Johnson S.D. 2010. Floral mimicry enhances pollen export: the evolution of pollination by sexual deceit outside of the Orchidaceae. The American Naturalist, 176, E143–E151.

Fan X.L., Barrett S.C.H., Lin H., Chen L.L., Gao J.Y. 2012. Rain pollination provides reproductive assurance in a deceptive orchid. Annals of Botany, 110(5), 953–958.

Gottsberger, G. 2012. How diverse are Annonaceae with regard to pollination? Botanical Journal of the Linnean Society, 169, 245–261.

Goss G.J. 1977. The reproductive biology of the epiphytic orchids of Florida. 6. *Polystachya flavescens* (Lindley) J.J. Smith. American Orchid Society Bulletin, 46, 990–994.

He M.G. 2010. Preliminary study on seed biology of orchid in Hainan. Master Dissertation. Graduate School of the Hainan University.

Li J.H., Liu Z.J., Salazar G.A., Bernhardt P., Perner H., Tomohisa Y., Jin X.H., Chung S.W., Luo Y.B. 2011. Molecular phylogeny of *Cypripedium* (Orchidaceae: Cypripedioideae) inferred from multiple nuclear and chloroplast regions. Molecular Phylogenetics and Evolution, 61, 308–320.

Lunau K. 2000. The ecology and evolution of visual pollen signals. Plant Systematics and Evolution, 222(1-4), 89–111.

Lunau K., Konzmann S., Winter L., Kamphausen V., Ren Z.X. 2017. Pollen and stamen mimicry: the alpine flora as a case study. Arthropod-Plant Interactions, 11(3), 1–21.

Lunau K., Ren Z.X., Fan X.Q., Trunschke J., Wang H. 2020. Nectar mimicry: a new phenomenon. Scientific Reports, 10, 7039.

Meve U., Liede S. 1994. Floral biology and pollination in stapeliads—new results and a literature review. Plant Systematics and Evolution, 192, 99–116.

Jersáková J., Johnson S.D., Kindlmann P. 2006. Mechanisms and evolution of deceptive pollination in orchids. Biological Reviews, 81(2), 219–235.

Jersáková J., Johnson S.D. 2006. Lack of floral nectar reduces self-pollination in a fly-pollinated orchid. Oecologia, 147, 60–68.

Jiang H., Kong J.J., Chen H.C., Xiang Z.Y., Zhang W.P., Han Z.D., Liao P.C., Lee Y.I. 2020. *Cypripedium subtropicum* (Orchidaceae) employs aphid colony mimicry to attract hoverfly (Syrphidae) pollinators. New phytologist, 227, 1213–1221.

Johnson S.D., Schiestl F.P. 2016. Floral mimicry. Oxford, UK: Oxford University Press.

Nilsson L.A. 1992. Orchid pollination biology. Trends in Ecology and Evolution, 7(8), 255–259.

Pansarin E.R., Amaral M.C.E. 2006. Biologia reprodutiva e polinizao de duas espécies de polystachya hook. no sudeste do brasil: evidência de pseudocleistogamia em polystachyeae (orchidaceae). Revista Brasilra De Botnica, 29(3), 423–432.

Pansarin E.R., Maciel A.A. 2017. Evolution of pollination systems involving edible trichomes in orchids. AoB PLANTS, 9(4).

Papiorek S., Junker R.R., Alves-dos-Santos I., Melo G.A.R., Amaral-Neto L.P., Sazima M., Wolowski M., Freitas L., Lunau K. 2016. Bees, birds and yellow flowers: pollinator-dependent convergent evolution of UV patterns. plant biology, 18(1), 46–55.

Ren Z.X., Li D.Z., Bernhardt P., Wang H. 2011. Flowers of *Cypripedium fargesii* (Orchidaceae) fool flat-footed flies (Platypezidae) by faking fungus-infected foliage. Proceedings of the National Academy of Sciences USA, 108, 7478–7480.

Sanguinetti A., Buzatto C.R., Pedron M., Davies K.L., de Abreu Ferreira P.M., Maldonado S., Singer R.B. 2012. Floral features, pollination biology and breeding system of Chloraea membranacea Lindl. (Orchidaceae: Chloraeinae). Annals of Botany, 110, 1607–1621.

Singer R.B., Flach A., Koehler S., Marsaioli A.J., Do Carmo E., Amaral M. 2004. Sexual mimicry in *Mormolyca ringens* (Lindl.) Schltr. (Orchidaceae: Maxillariinae). Annals of Botany, 93, 755–762.

Singer R.B., Koehler S. 2004. Pollinarium morphology and floral rewards in Brazilian Maxillariinae (Orchidaceae). Annals of Botany, 93, 39–51.

Van Waes J.M., Debergh P.C. 1986. Adaptation of the tetrazolium method for testing the seed viability, and scanning electron microscopy study of some Western European orchids. Physiologia plantarum, 66(3), 435–442.

Wang Y.Q., Zhang D.X., Renner S.S., Chen Z.Y. 2004. A new self-pollination mechanism. Nature, 431(7004), 39–40.

Wester P., Lunau K. 2017. Plant-pollinator communication. Adv Bot Res 82:225–257.

Zheng G.L., Li P. (2009) Study on plant resources and breeding system of *Cypripedium* in Sichuan. Journal of Anhui Agricultural Sciences, 37, 5468–5469. (in Chinese with English abstract).

